# Assessing the role of collagen to improve understanding of disease pathophysiology in patients with acute recurrent tonsillitis

**DOI:** 10.1101/2025.07.07.663453

**Authors:** Kay Polland, Megan Clapperton, Thushitha Kunanandam, Catalina D Florea, Catriona M Douglas, Gail McConnell

## Abstract

Acute recurrent tonsillitis (ART) and obstructive sleep apnoea (OSA) are the most common indications for tonsillectomy worldwide however the disease pathophysiology is poorly understood. Acute recurrent tonsillitis causes repeated episodes of inflammation within the tonsils. Repeated episodes of inflammation can lead to scarring of tissue due to an overproduction of extracellular matrix proteins such as collagen, which in return can cause decreased immunological function. In this study we assess collagen prevalence within ART and OSA tonsils with the aim of better understanding the underlying disease pathophysiology. By performing label-free second harmonic generation (SHG) imaging of tonsil tissue from patients with ART and OSA it is possible to enable measurement of the relative abundance of immature and mature type I collagen within *ex vivo* paediatric palatine tonsils. We have demonstrated that there is a significant difference in the mean area fraction of immature type I collagen with a higher relative fraction in patients with ART compared to those with OSA (p = 0.0059). However, there was no statistically significant difference in the mean area fraction of mature type I collagen in patients with ART and OSA (p = 0.61). Spatial analysis demonstrated that there was a significantly fraction in tonsillar tissue from patients with ART of immature type I collagen in the epithelia dominant regions (p = 0.0074) and the interior regions (p = 0.0023). Our results provide new information regarding these two common paediatric diseases, giving us an improved insight into the disease pathophysiology.

## 1. Introduction

Tonsillar diseases are amongst the most common illnesses GPs will encounter, with acute recurrent tonsillitis (ART) being particularly prevalent between the ages of 5-15 years old (Georgalas et al. 2009). The most common signs and symptoms of ART include throat pain, fever, odynophagia and inflammation of the tonsils. This causes a significant impact on a patient’s quality of life, requiring time off education and social activities. Children with acute tonsillitis often require repeated courses of antibiotics to treat infection, with concerns arising around antimicrobial resistance (Pelucchi et al. 2012). If patients fulfill the SIGN 7,5,3 rule, they are referred for consideration of tonsillectomy (SIGN 2010). Obstructive sleep apnoea (OSA) is the complete or partial collapse of the upper airway due to adenotonsillar hypertrophy, with a high prevalence in children between 2-8 years of age (Garg et al. 2017)(Li et al. 2016)(Schwengel et al. 2014). Children with OSA present with sleep disturbance which can lead to excessive daytime sleepiness, snoring, morning headaches, and poor focus during the day. Definitive treatment of OSA is with tonsillectomy. Tonsillectomy is the most common operation performed in children, however, it is not without morbidity; up to 20% of tonsillectomy patients get readmitted to hospital in the two-week post-operative period due to pain and bleeding.

Collagen is the most abundant protein within the body, providing structure and support for connective tissue, bones, muscles, and skin within the body. Type I collagen is the most abundant collagen within the body and forms the extracellular matrix (ECM) within tonsillar tissue. The primary role of collagen is to manage the health of connective tissue and the mechanical structures of our skin. It is essential for tissue restructuring, blood clotting and helps cells to adhere to one another by binding to their triple helices. Immature collagen fibrils are less well organised and have a lower degree of cross-linking, making them less stable and increasing the risk of degradation. Immature collagen is involved in early wound healing stages and if unregulated it can lead to significant scarring and chronic wounds (Wang et al. 2018). Mature type I collagen has a higher degree of crosslinking providing structure and strength. Tissue damage instigates inflammation and degradation of the ECM and whilst collagen rebuilds the damaged ECM it can become uncontrollable and lead to abnormal growth (Singh et al. 2023). This abnormal growth in the ECM can promote disease development, such as fibrosis, (Urbanczyk et al. 2019) and can play a vital role in various biological processes of the body.

Previous work has used absorption staining methods to assess the abundance of type I collagen in tonsillar disease using a 400x magnification. However, this method produces images with a very small field of view and hence it is difficult to provide statistically robust sampled data that are representative of the disease (Dantas 2012). Additionally, absorption staining methods for labelling of collagen are subject to incomplete and non-specific binding, which can lead to bias in the data and interpretation of results (Buchwalow et al. 2011).

Second harmonic generation (SHG) microscopy is a nonlinear optical technique in which two photons, usually of near-infrared light, interact with a non-centrosymmetric material to produce a coherent beam of light at exactly half the wavelength of the input radiation (Aghigh et al. 2023). SHG imaging has been applied to the study of fibrosis and neck squamous cell carcinoma, and previous work has shown that SHG imaging of immature type I collagen and mature type I collagen can be performed on a single microscope by collection of the forward and backwards SHG signals respectively (Williams et al. 2005) (Chen et al. 2012). Importantly, SHG imaging does not require a label to produce contrast, therefore there is no risk of photobleaching that can complicate observation and quantification. Using label-free SHG imaging, we aimed to assess the relative abundance of immature type I collagen and mature type I collagen, in patients with either ART or OSA. Furthermore, we aimed to assess whether the abundance of immature and mature collagen in these two patient groups was dependent on the anatomical location within the tonsil.

## 2. Methods

### 2.1 Specimen Preparation

Tonsil samples (n = 15) were collected from the Royal Hospital for Children, Glasgow, UK. The ethics were approved by Biorepository Management Committee of NHS Greater Glasgow and Clyde, UK (NHS Biorep 548). Specimens were blinded to prevent bias, with a total of 7 tonsils from patients with OSA and 8 tonsils from patients with ART used in this study. Specimens were then rapidly transported to the Strathclyde Institute for Pharmacy and Biomedical Sciences at the University of Strathclyde in a sterile saline solution (0.9% Sodium chloride, Baxter Healthcare Ltd, UK). Whole fresh tonsils were frozen in optical cutting temperature (OCT) embedding matrix for frozen tissue (KMA-0100-00A, Cell Path, UK) then were sectioned to a thickness of 20 µm using a microtome (CM1950, Leica Microsystems, Germany) and specimens were mounted on microscope slides (Z692255, Sigma-Aldrich, USA). This thickness was chosen to minimise tissue folding and tearing. Tissue sections were fixed in 4% paraformaldehyde followed by a series of 3 washes for 5 minutes each using 1X phosphate buffered saline (PBS) (10010023, ThermoFisher Scientific, USA). The specimens were mounted under Type 1.5 coverslips using VectaMount (H-5000, Vector Laboratories), and allowed to set overnight before imaging.

### 2.2 SHG microscope configuration

The imaging set-up is shown in Figure 1(a). Images were acquired using a laser scanning microscope (SP5, Leica) equipped with a horizontally polarised, ultrashort pulsed Ti:Sapphire laser source (Chameleon Vision II, Coherent) tuned to a wavelength of 850 nm. This laser source had an 8 nm full-width at half-maximum spectral bandwidth and produced pulses of 140 fs duration at a repetition rate of 80 MHz. This source was used for second-harmonic generation of collagen, with the laser parameters chosen based on previous work reporting imaging of type I collagen in sections of mouse skin (Campagnola et al. 2002).

**Figure 1.**
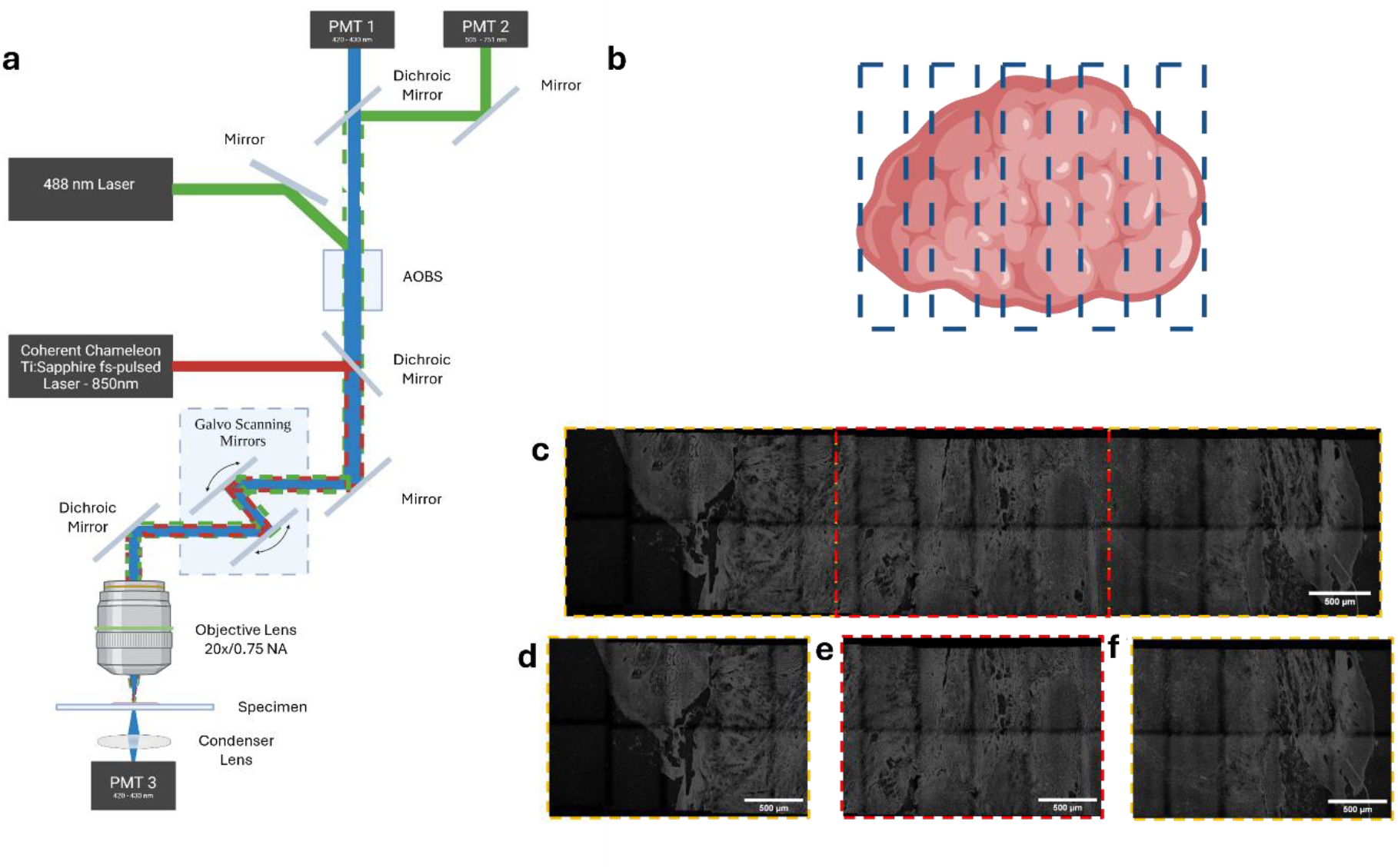
(a) Schematic of the Leica SP5 imaging system set for multimodal imaging. A horizontally polarised ultrashort pulsed Coherent Chameleon Ti-Sapphire Vision II laser source was tuned to a wavelength of 850 nm and was used for second harmonic generation of the collagen. A separate 488 nm laser was used to excite tissue autofluorescence. Forward and backward second harmonic generation (SHG) signals, corresponding to mature type I collagen and immature type I collagen respectively, were each detected between 422nm to 427 nm (PMT 3 and PMT 1 respectively), and autofluorescence of the tissue was detected between 505-751 nm (PMT 2). (b) Schematic diagram showing the relative position of the imaged ROIs in the tonsil sections. (c) Stitched and tiled SHG image dataset taken across the width of the tonsil. Yellow and red boxes highlight sub-regions of interest (ROIs). (d), (e) and (f) are regions of (c), with (d) and (f) showing the position of the epithelial dominant tonsil and (e) showing the tonsil interior region.

A separate 488 nm Argon laser was used to excite tissue autofluorescence for landmarking and calculating the tissue area. Three detection channels were used, recording the backward SHG (PMT 1), tissue autofluorescence (PMT 2), and the forward SHG (PMT 3) signals respectively. The detection wavelengths for PMT 1 were 422 nm to 427 nm, while PMT 2 captured signals from wavelengths of 505 nm to 750 nm and PMT 3 recorded information from wavelengths of 422nm to 427 nm. A 20x/0.75 numerical aperture air immersion lens was used. This provided imaging of individual regions of the tissue preparation of 500 µm in diameter. However, the tonsil tissue sections exceeded several millimetres in diameter, and hence image stitching and tiling was used to cover larger regions.

Individual images were captured at a rate of 400 lines per second and with 512 × 512 pixels averaged over 3 frames to improve signal to noise ratio, with a time per frame of 1.2 seconds, without averaging. While this did not meet the Nyquist sampling criterion (Landau 1967) for resolution, this number of pixels and scan speed was chosen to produce images of sub-cellular resolution at high speeds. Auto-stitching was chosen in the Leica SP5 control software (LAS-AF, Leica) to produce high-quality mosaics of individual image tiles. A 10% overlap in the X and Y directions was applied. This was performed to increase the imaging area to improve cell statistics. Each tonsil section was imaged, with 5 regions of interest (ROIs). The minimum ROI was 1436 µm x 3177 µm and the maximum ROI was 3133 µm x 9301 µm, as shown in Figure 1(b), resulting in a total of 40 ROIs for ART tissue and 35 ROIs for OSA tissue. Images were saved in the proprietary Leica.lif format, and data were exported as 8-bit .TIFF files for presentation and analysis. False colours were applied in Fiji (Schindelin et al. 2012).

### 2.3 Image processing and image data analysis

Image analysis of raw 8-bit data was performed in Fiji (Schindelin et al. 2012) to assesss the location and abundance of immature type I collagen and mature type I collagen across the tonsils from patients with ART and OSA. A rolling ball radius was applied to channel 3 images and default thresholding (Sternberg 1983) of each image ROI for all channels was obtained and the areas of the immature type I collagen, mature type I collagen, and tissue autofluorescence were measured. Tissue autofluorescence was normalised to 100% of the area and the area fraction of immature type I collagen was calculated by taking the normalisation multiplier for the tissue autofluorescence area fraction and multiplying by the immature type I collagen value. A similar operation was performed for mature type I collagen to calculate the area fraction of mature type I collagen. The mean area fractions (MAFs) for both collagen types in both ART and OSA tissues were then calculated by taking the mean of the individual area fractions from the ROIs. Images for presentation purposes were contrast adjusted using CLAHE with the default parameters (Zuiderveld 1994), but all analysis was performed prior to adjustment using CLAHE. The ROUT outlier removal test (GraphPad Prism v.8.0.2, GraphPad Software) with Q = 1%, was performed (Motulsky et al. 2006). Mean values of the MAF for both immature type I and mature type I collagen were determined from the processed datasets along with the standard deviation of the dataset. All data sets were non-normally distributed, therefore a Mann-Whitney U Test was performed to determine the significance of data in the two patient cohorts. Plots were generated in GraphPad Prism.

Further analysis was performed to determine the spatial location of immature type I collagen across patient cohorts. ROIs were splt into three sub-regions of interest, with two regions containing tonsil epithelium and one with the tonsil interior. This is shown schematically in Figure 1(c). The method described above to calculate the MAF of the ROIs was then applied to these sub-regions to reveal if the immature type I collagen burden was spatially localised or spread globally throughout the tonsil. ROIs which contained only epithelial edge were not assessed as they did not contain a well defined interior.

## 3. Results

### 3.1 SHG imaging reveals differences in type I collagen abundance and location in tonsillar tissue from patients with ART and patients with OSA

Representative images of tonsil sections from patients with ART are shown in Figure 2. Figures 2(a) and 2(b) show tissue autofluorescence, 2(c) and 2(d) show type I mature collagen, 2(e) and 2(f) show type I immature collagen, and 2(g) and 2(h) are false colour merges of (a), (c), and (g) and (b), (d), and (f) respectively. Stitching and tiling artefacts are visible in each of the image datasets. These images show morphological differences in the amount of collagen types. From a visual assessment, both type I immature and mature collagen have a greater abundance in tissue from patients with ART compared to that from patients with OSA. Spatially, both immature type I and mature type I collagen were located close to the epithelium in both disease types. In addition, collagen can be seen in the interior of the tonsils in patients with ART while the same collagen presence is not visible in the tissue from patients with OSA.

**Figure 2.**
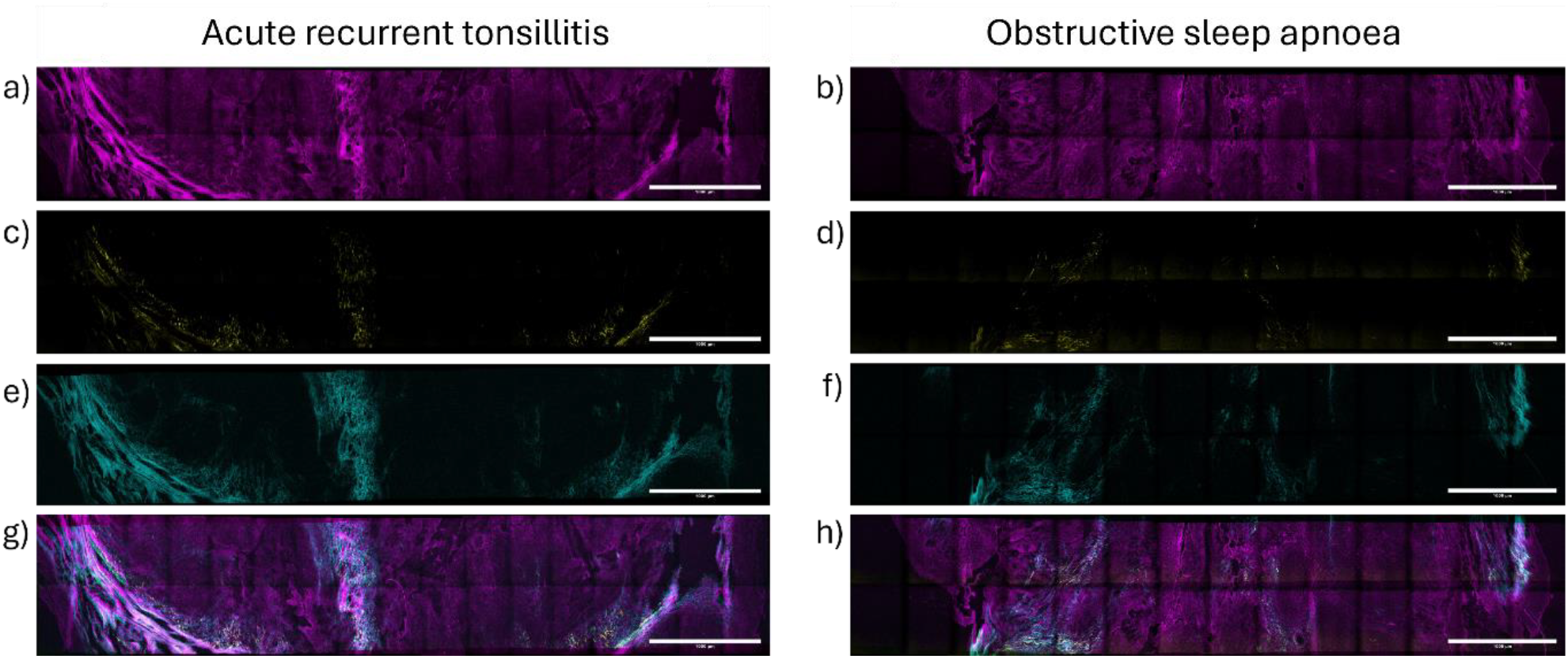
Imaged regions of interest in tonsil tissue from patients with ART and OSA. (a) and (b) show images of tissue autofluorescence (magenta) from OSA and ART respectively. SHG images of type I mature collagen (yellow) distribution in tissue from patients with ART and OSA are shown in (c) and (d), while SHG images of type I immature collagen (cyan) are shown in (e) and (f) for the same diseases. In (g) and (h), false-colour merge images of (a), (c), and (e), and (b), (d) and (f) respectively are presented, showing the location of type I mature and type I immature collagen in the tonsil tissue. These images have been adjusted using CLAHE within Fiji for presentation purposes (Schindelin et al. 2012)(Zuiderveld 1994). Scale bars = 1 mm in all images.

### 3.2 The MAF of immature I collagen in tonsillar tissue differs in patients with ART and patients with OSA

Figure 3 shows the results of the MAF analysis of immature type I and mature type I collagen in n = 75 tonsil ROIs. A a significantly higher MAF of immature type I collagen was observed in patients with ART (mean value = 8.79, n = 33 ROIs, Outliers removed = 2) compared to patients with OSA (mean value = 3.45, n = 25 ROIs, Outliers removed = 5), p = 0.0059. There was a higher MAF of mature type I collagen in patients with ART (mean value = 1.37, n = 32 ROIs, Outliers = 3) than those with OSA (mean value = 1.16, n = 30 ROIs, Outliers removed = 0), but this measurement was not significant (p = 0.61).

**Figure 3.**
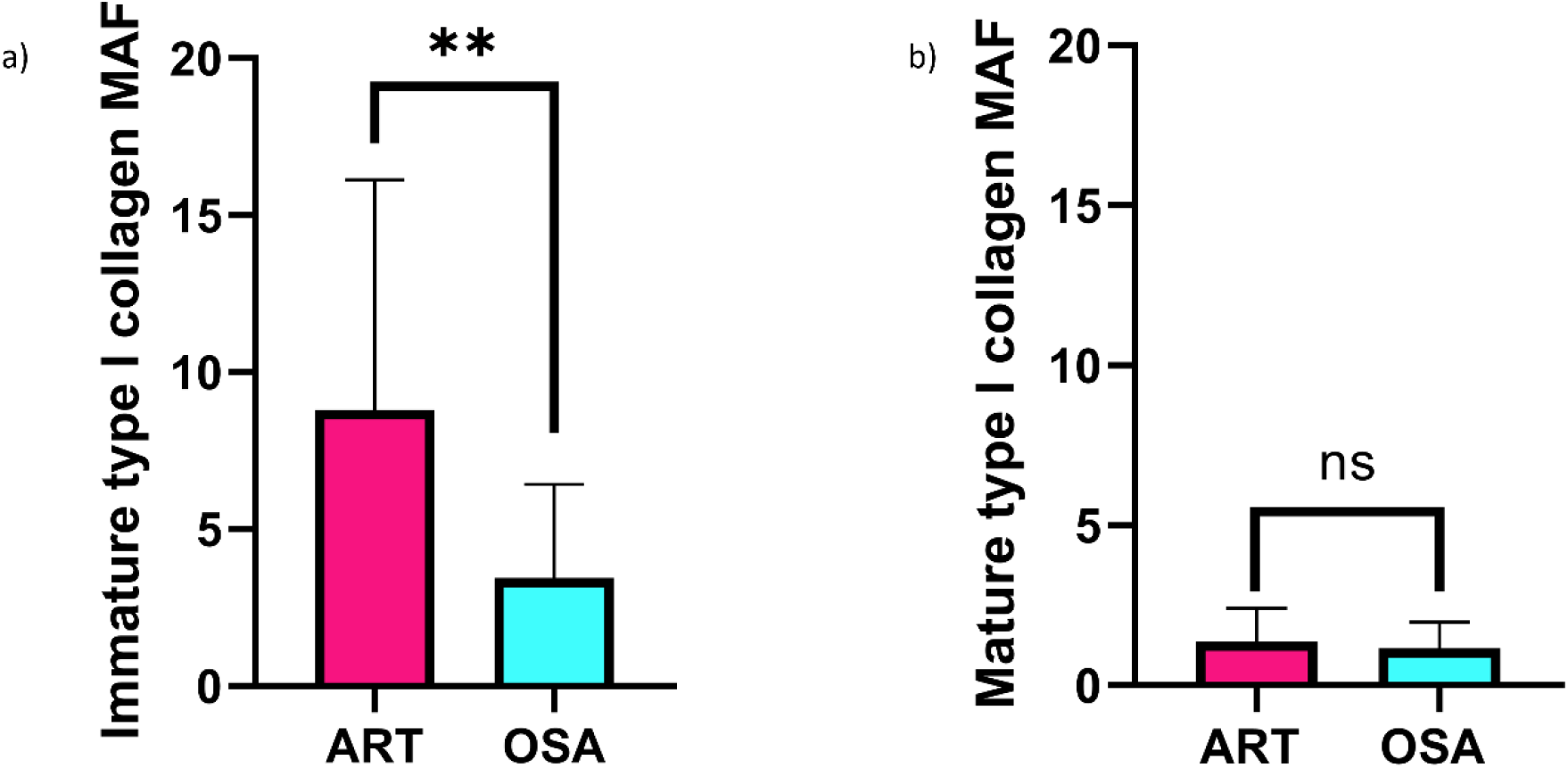
Mean area fraction (MAF) measurements for tonsil tissue sections for patients with OSA (cyan) and ART (magenta). (a) The MAF of mature type I collagen is higher in tissue from patients with ART patients than in tissue from patients with OSA but this difference is not statistically significant (p = 0.61). (b) The MAF of immature type I immature collagen is significantly higher in tissue from patients with ART when compared with the same measurement in tissue from patients with OSA (p = 0.0059).

### 3.3 The MAF of immature type I collagen differs in patients with ART and OSA at the tonsil epithelium but not in the tonsil interior

Figure 4 shows the measured MAFs of immature type I collagen in epithelial dominant regions and interior regions of the tonsil tissue from patients with ART and OSA respectively. There was a statistically significant increase in immature type I collagen in the tonsil epithelial dominant regions of tissue from patients with ART (mean value = 8.45, n = 51, Outliers removed = 3) compared with tonsil tissue from patients with OSA (mean value = 2.79, n = 40, Outliers removed = 10), p < 0.0001. However, there was no significance in the MAF of immature type I collagen in the interior of tissue from patients withART (mean value = 5.72, n = 24, Outliers removed = 3) when compared to tonsil tissue from patients with OSA (mean value = 2.59, n = 21, Outliers removed = 4), p = 0.16.

**Figure 4.**
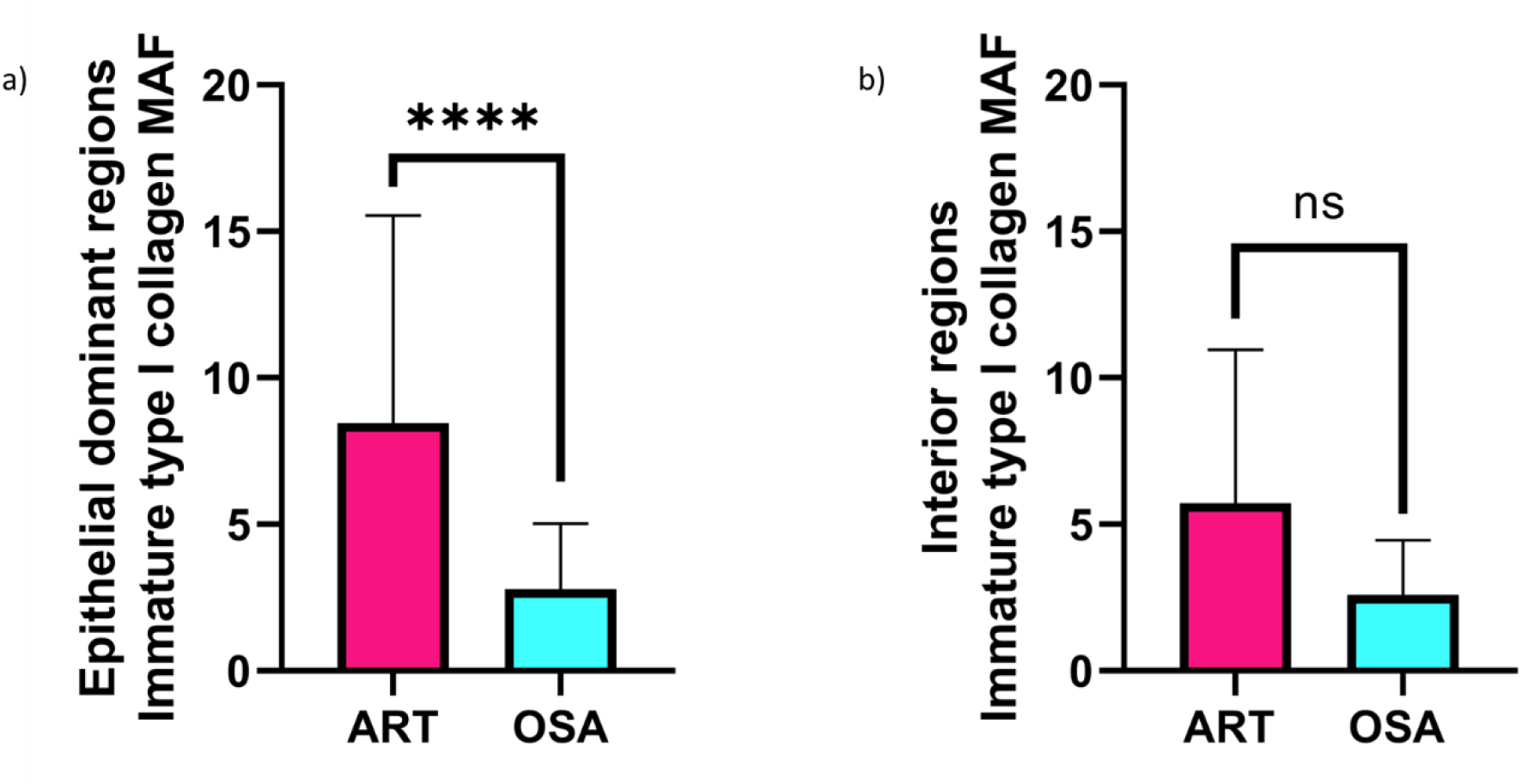
Mean area fraction (MAF) for tonsil sections from patients with OSA (cyan) and ART (magenta) in epithelial dominant regions (a) where there is highly statistically significant increase in immature type I collagen in patients with ART than those with OSA (p < 0.0001), and the interior regions (b) of the tonsil where there was no statistical significance in immature type I collagen when comparing between patients with ART and OSA (p = 0.16).

## 4. Discussion

The purpose of this study was to assess the relative abundance of mature type I collagen and immature type I collagen in patients with 2 common tonsillar diseases, with a view to better understanding the potential role of type I collagen in disease biology. Through the use of SHG imaging and image analysis, our findings revealed a significantly higher mean area fraction of immature type I collagen across both the tonsil epithelial dominant regions and interior regions in tissue from patients with ART compared to OSA patient tissue.

The observed differences in collagen maturity suggest a potential pathophysiological role for immature type I collagen in ART. Type I collagen is a fundamental structural protein in the extracellular matrix (ECM), and its remodeling is a hallmark of fibrosis and chronic inflammatory diseases (Herrera et al. 2018). High levels of immature type I collagen fibrils have been implicated as early indicators in the pathogenesis of fibrotic diseases such as liver, pulmonary, and kidney fibrosis (Jones 2018; Dooling 2023). In contrast, elevated levels of mature type I collagen have been associated with scleroderma, where excessive collagen cross-linking leads to tissue stiffening and dysfunction (Alghamdi et al. 2024). Our findings suggest that an imbalance in collagen maturation may contribute to the recurrent inflammatory processes in ART, leading to persistent tissue remodeling and fibrosis.

This study demonstrates a significant difference in the mean abundance of immature type I collagen between patients with ART and OSA (p = 0.0059), with ART patients exhibiting significantly higher levels. This suggests a potential involvement of immature type I collagen in the pathophysiology of ART. The increased presence of immature collagen may indicate a failure in normal tissue remodeling processes, resulting in a predisposition to chronic inflammation and fibrosis. Future research could investigate whether antifibrotic treatments, currently used for conditions such as idiopathic pulmonary fibrosis (Maher and Strek 2019), may be beneficial in treating recurrent tonsillitis, thereby reducing the clinical burden on healthcare systems and improving patient outcomes by potentially avoiding invasive tonsillectomy.

Collagen plays a vital role in tissue repair and immune regulation within the palatine tonsils. The tonsils are a secondary lymphoid organ, constantly exposed to pathogens, making them a site of dynamic immune responses and tissue remodeling. The interplay between collagen production and degradation is crucial in maintaining tonsillar homeostasis. Fibroblasts and myofibroblasts are responsible for ECM turnover, producing collagen to support tissue integrity while responding to inflammatory stimuli (Li et al. 2022)(Otranto et al. 2012). In chronic inflammatory conditions such as ART, dysregulated fibroblast activity may lead to an excessive accumulation of immature collagen, impairing normal tissue function and contributing to recurrent infection and fibrosis.

Our results are likely to be of significant immunological interest and importance. For example, it is has been shown that IL-10 is expressed in the palatine tonsil, and that the average expression of IL-10 depends on the extent of tonsillar hypertrophy (Mikola etal. 2018). Recent reports suggest that IL-10 regulates collagen and collagenase gene expression in fibroblasts, and hence further research could investigate if there is a correlation between IL-10 and immature collagen production in patients with ART. It has also been reported that IL-37, a fundamental inhibitor of innate immunity, is slightly elevated in patients with tonsillar hypertrophy compared to patients with ART (Mikola et al. 2018). IL-37 is mainly expressed in tonsillar B cells (Jiang et al. 2023) and in lung has been shown to attenuate fibrosis by inducing autophagy (Kim et al. 2019). An interesting area of possible study would be to understand whether the antifibrotic activity of IL-37 could be used as a therapeutic target in tonsillar diseases. Furthermore as Il-1 is known to be a regulator of fibroblasts inflammatory responses and synthesis of collagen in airways, as well as fibronectin (Osei et al. 2020), it could be a further area of research to study this alongside collagen abundance to assess whether a lack of Il-1 is leading to low ability to repair and remodel tissue.

Cytokine-driven regulation of collagen synthesis in the tonsils is another critical aspect of disease pathogenesis. Pro-inflammatory cytokines, including transforming growth factor-beta (TGF-β), tumor necrosis factor-alpha (TNF-α), and interleukins such as IL-6 and IL-17, are known to influence fibroblast activity and ECM remodeling (Johnson et al. 2020). TGF-β, a potent fibrogenic cytokine, has been shown to drive excessive collagen deposition in fibrotic diseases and may play a similar role in ART (Kim et al. 2018). Conversely, anti-inflammatory cytokines such as IL-10 can suppress collagen synthesis and regulate fibroblast proliferation, potentially mitigating fibrosis in tonsillar tissues.

Furthermore, recent studies (Cabral-Pacheco et al. 2020) highlight the role of matrix metalloproteinases (MMPs) in ECM remodeling. MMPs, particularly MMP-1 and MMP-9, are responsible for collagen degradation and remodeling. Dysregulated MMP activity has been linked to various fibrotic conditions and may contribute to the excessive accumulation of immature collagen in ART (Giannandrea and Parks 2014). Evaluating the balance between collagen MAF using SHG and degradation through MMP and tissue inhibitor of metalloproteinase (TIMP) expression could provide valuable insights into the pathophysiology of ART.

From a clinical perspective, these findings have important implications for treatment strategies. Patients with ART frequently undergo tonsillectomy due to recurrent infections and tissue hypertrophy. Understanding the role of collagen in disease progression could open avenues for less invasive therapeutic interventions. For instance, targeting collagen maturation pathways with antifibrotic agents such as pirfenidone or nintedanib, both of which are approved for pulmonary fibrosis (Bahudhanapati et al. 2017)(NICE 2023), could be explored as potential topical treatments for ART.

Another area of interest is the role of the microbiome in modulating ECM remodeling. The tonsillar crypts harbor diverse microbial communities that interact with host immune responses. Emerging evidence suggests that dysbiosis may contribute to chronic inflammation and fibrosis (De Minicis et al. 2014)(Loomba et al. 2017). Investigating the interplay between the tonsillar microbiome, immune regulation, and collagen remodeling using SHG imaging could provide novel insights into the etiology of ART and potential microbiome-targeted therapies.

Given that SHG signal from immature type I collagen is collected in the backwards direction, unlike the SHG signal from mature type I collagen, which is detected in the forwards direction, our data suggest that it may be possible to use endoscopic illumination and detection for SHG imaging of the human tonsil *in situ* for the purpose of diagnosis of ART. SHG endoscopy and microendoscopy have been used previously in mouse, including for the study of intestinal microstructure (Gu et al. 2014) and in humans, including applications in the visualisation of cervical remodelling (Zhang et al. 2012). These existing optical technologies could be repurposed or adapted for the study of immature type I collagen in situ in the human tonsil. This could allow for possible assessment of severity in tonsillar diseases. Importantly, with a statistically significant difference in SHG signal from immature type I collagen measured irrespective of the location in patients with ART, further in-vivo studies could be done without imaging of a pre-specified or well-defined area of the tonsil. Severity of disease alongside collagen abundance could be studied to investigate the impact of collagen on tonsillar diseases using this technology. Prior to embarking on endoscopic imaging, however, it would be useful to perform backwards-SHG imaging of whole mounts of tonsil tissue to ensure that our results were replicated in ultra-thick tissue sections, and the same trends were observed as in the thin tissue sections we have studied thus far.

By integrating optical physics with biological and clinical perspectives, this study lays the groundwork for further exploration into the mechanisms driving tonsillar disease and potential therapeutic targets. The ability to quantitatively assess collagen MAF through SHG imaging presents a promising avenue for improving diagnosis and treatment of ART, potentially reducing the need for surgical interventions and improving patient quality of life.

## 5. Conclusion

We have used label-free multimodal microscopy to visualise the location and image analysis to measure the relative abundance of type I mature and type I immature collagen in tonsils from patients with either ART or OSA. We observed a statistically significant higher relative abundance of type I immature collagen in patients with ART compared to those with OSA. This provides a greater understanding of the disease pathophysiology, which would allow for further work in developing minimally invasive diagnostic tools for tonsillar diseases, such as the design of an endoscope for multimodal *in vivo* imaging. Additonally, this increased understanding of collagen abundance and distribution could be explored alongside biomarkers for relevant interleukins to provide more detailed insights into disease pathology.

## Acknowledgements

This work was supported by a studentship to K.P. from Medical Research Scotland (PhD-50367-2021) and CoolLED Ltd. M.C. was supported by the Glasgow Children’s Hospital Charity, grant number GCHCRF/PHD/2020/02. C.M.D. was partly supported by the Chief Scientific Office and UK Research and Innovation, grant number MR/W030381/1. G.M. was funded by UK Research and Innovation, grant numbers BB/T011602/1, BB/X005178/1, BB/V019643/1, MR/K015583/1, and The Leverhulme Trust.

## Notes

### Competing Interest Statement

The authors have declared no competing interest.

